# Investigating trait variability of gene co-expression network architecture in brain by manipulating genomic signatures of schizophrenia risk

**DOI:** 10.1101/2021.05.04.442668

**Authors:** Eugenia Radulescu, Qiang Chen, Giulio Pergola, Nicholas J Eagles, Joshua M Stolz, Joo Heon Shin, Thomas M Hyde, Joel E Kleinman, Daniel R Weinberger

**Affiliations:** Lieber Institute for Brain Development, Johns Hopkins Medical Campus, Baltimore, MD, 21205, USA; Group of Psychiatric Neuroscience, Department of Basic Medical Sciences, Neuroscience and Sense Organs, University of Bari Aldo Moro, Bari, Italy; Department of Neurology, Johns Hopkins School of Medicine, Baltimore, MD 21205, USA; Department of Neuroscience, Johns Hopkins School of Medicine, Baltimore, MD 21205, USA; Department of Psychiatry and Behavioral Sciences, Johns Hopkins School of Medicine, Baltimore, MD 21205, USA; McKusick-Nathans Department of Genetic Medicine, Johns Hopkins School of Medicine, Baltimore, MD 21205, USA

## Abstract

While the role of genomic risk for schizophrenia on brain gene co-expression networks has been described, the patterns of its manifestations are varied and complex. To acquire a deeper understanding of this issue, we implemented a novel approach to network construction by manipulating the RNA-Seq expression input to “integrate” or “remove” the “modulatory” effects of genomic risk for schizophrenia. We created co-expression networks in DLPFC from the adjusted expression input and compared them in terms of gene overlap and connectivity. We used linear regression models to remove variance explained by RNA quality, cell type proportion, age, sex and genetic ancestry. We also created co-expression networks based on the genomic profile of a normative trait, height, as a “negative control”; we also applied the same analytical approach in two independent samples: LIBD Human Brain Repository (HBR) (N=78 brains, European ancestry) and Common Mind Consortium (CMC) (N=116 brains, European ancestry). In addition to direct comparisons, we explored the biological plausibility of the differential gene clusters between co-expression networks by testing them for enrichment in relevant gene ontologies and gene sets of interest (PGC2-CLOZUK GWAS significant loci genes, height GWAS significant loci genes, genes in synaptic ontologies-SynGO and genes of the “druggable genome”). We identify several key aspects of the role of genomic risk for schizophrenia in brain co-expression networks: 1) Variability of co-expression modules with “integration” or “removal” of genomic profiles of complex traits (normal or pathological); 2) Biological plausibility of gene sets represented in the differential co-expression contrasts and potential relevance for illness etiopathogenesis; 3) Non-preferential mapping of schizophrenia GWAS loci genes to network areas apparently influenced by the genomic risk score. Overall, our study supports the notion that genomic risk for schizophrenia has an extensive and non-linear effect on brain gene co-expression networks that possibly manifests as a molecular background for gene-gene, gene-environment interactions that affect various biological pathways.

## INTRODUCTION

Crystalized in the name of the most severe psychotic illness - schizophrenia (SCZ) - is the definitory feature of this condition, a failure of connectedness with profound consequences for a person’s individual and social life. The fundamental notion - first proffered by Eugen Bleuler in his use of the term ‘schizophrenia’[1] - that thoughts and ideas are disconnected, has been abstracted at various biological levels, including macrocircuit level in neuroimaging connectivity studies [2-4] and microcircuit level in gene co-expression studies in post mortem brain. These approaches have been exploited as a foundation for understanding brain dysfunction in this disorder and its molecular origins [5-12].

Gene co-expression network analysis represented by one standard application, Weighted Gene Co-expression Network Analysis (WGCNA), is a powerful tool for highlighting subjacent biological mechanisms associated with risk for neuropsychiatric disorders [13]. Previous studies with WGCNA on RNA-Seq data from postmortem DLPFC of affected and non-affected individuals found various degrees of association between genetic liability for SCZ and gene co-expression networks [5, 9-10, 12, 14-15]. Genetic associations have been used to prioritize clusters/modules of co-expressed genes with a link to risk for SCZ [5].

Notwithstanding the potential of gene co-expression analysis, however, this approach is not without important caveats. With its correlational nature, WGCNA inherently provides an output that represents an indirect association with a trait of interest and it, therefore, requires independent evidence for validation of results [16]. Furthermore, like any linear representation of complex traits with most likely non-linear architectures of poorly understood interactions of genes, co-expression network construction is sensitive to many biological factors (e.g. age, cell composition) and technical artifacts (e.g. degradation) [17-19].

While many of these factors in network construction are potential confounders or limitations, they might also be considered opportunities to systematically manipulate networks to extract novel and biologically plausible information about gene co-expression association with complex traits. For example, the variability in network construction related to diverse factors that influence gene expression might be intentionally utilized to leverage the effect of specific factors of interest and to remove the effects of factors not of particular interest (e.g. technical artifacts or depending on the question, genetic risk), and thus “relatively isolate” specific effects.

In the present study, we took this novel approach to explore the impact on co-expression network architecture of varying gene expression input based on genetic risk. Among the possible drivers of RNA measures from bulk tissue co-expression networks are many biological factors in addition to genome profiles, including cell type composition [17], age [18], neuropathological factors (i.e., disease state [7-8]), as well as potential technical artifacts (i.e., RNA quality [19]). Here we propose that individual genomic profiles of complex traits will have distinct signatures on the co-expression network architecture that can be “relatively isolated” and this property can be used to contrast the contribution of genomic profiles of psychopathological versus normal traits. Specifically, we hypothesize that we can “extract” the influence of genomic risk for schizophrenia on co-expression network configuration from background network configuration. To test this hypothesis, we use statistical models of regression to adjust the input for co-expression network re-construction in a way that alternatively integrates or removes the potential effects of genomic profiles on gene expression. We then use the adjusted expression inputs to re-construct co-expression networks and compare them directly.

While the previous WGCNA studies addressing schizophrenia were designed to compare postmortem brain co-expression networks of patients and those of neurotypical individuals, to the best of our knowledge, no study described the architecture of the “generic background” co-expression network and its zones that are vulnerable to genetic risk for SCZ. To obviate the potential confounders of the state of illness and associated treatment, we focus these analyses on the DLPFC from neurotypical adult brain. In this study, we have specifically tested for evidence of genomic modulatory effects on brain co-expression networks by our trait of interest, i.e. genomic risk for schizophrenia (SCZ), and we use a normative trait, height (Ht), as a contrast for internal validation (“negative control”) and external comparator. Moreover, we apply the same methodological strategy to two independent gene expression datasets with the purpose of replication and external validation.

## METHODS

### Postmortem brain samples and RNA-Seq processing

Data was acquired from assays of postmortem human brain tissue from the LIBD Human Brain Repository (HBR), collected under a protocol of standardized brain acquisition, processing, and curation (location, legal authorizations, informed consent, clinical review/ diagnosis) described elsewhere [4, 20]. The RNA pre-processing pipeline and tissue quality check also have been detailed previously [20].

As an independent replication sample, we used the Common Mind Consortium (CMC) DLPFC RNA-Seq data (release 1) processed with SPEAQeasy (methodological details about brain collection and RNA-Seq treatment in **supplementary methods-SM1**; of note, for space economy and organization purposes, we further label supplementary methods by the numbered acronym **SM)**.

After pre-processing, two final gene expression data sets from LIBD (78 DLPFC samples) and CMC (116 DLPFC samples) from neurotypical adults of European ancestry were retained for further analysis (demographic characteristics in **figure 1A**).

**Figure 1:**
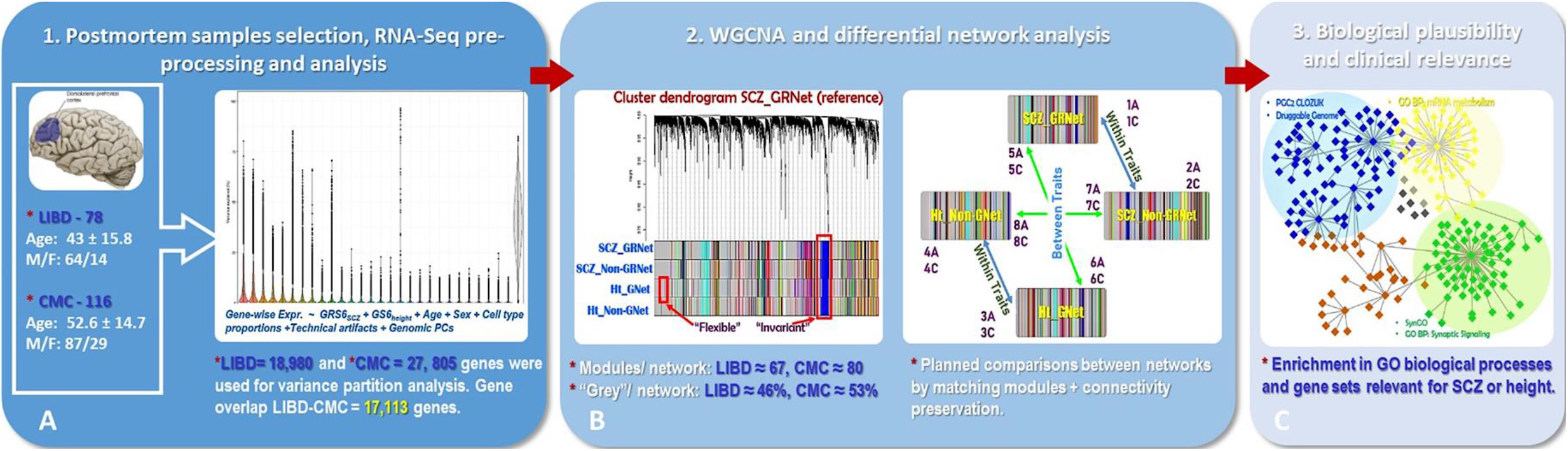
Analytical pipeline. ***A*:** Left panel-Demographic characteristics of the LIBD-CMC datasets; LIBD and CMC datasets significantly differ by age (Kruskal-Wallis chi-squared=16.118, df=1, p=5.951e-05), but not by gender (Pearson’s chi-squared=0.96649, p=0.326) (significance is calculated with Kruskal-Wallis test, respectively Pearson’s chi-squared test (for details see **supplementary material**); right panel: genome-wide violin plot that exemplifies the distribution of variance explained by each regressor across all genes. ***B:*** Left panel-Cluster dendrogram showing the correspondence between modules of a reference network and matched modules from the other three networks; modules from one network with no counterpart in the other three networks by gene overlap are part of the “flexible” sub-networks; conversely, modules of one network with correspondent in all other three networks are part of “invariant” sub-networks if gene overlap is greater than 20 genes. Right panel-Schematic representation of the 16 contrasts run for differential network analysis and characterization of the “flexible” sub-networks: 1A-4A = “Within traits” (WT) contrasts by matching modules from SCZ networks (Con 1A-2A: SCZ_GRNet vs. SCZ_nonGRNet and reverse), respectively “height” networks (Con 3A-4A: Ht_GNet vs. Ht_nonGNet and reverse); 5A-8A = “Between traits” (BT) contrasts by matching SCZ and “height” networks (Con 5A-6A: SCZ_GRNet vs. Ht_GNet and reverse; Con 7A-8A: SCZ_nonGRNet vs. Ht_nonGNet and reverse). Contrasts 1C-8C are connectivity based contrasts following the same directionality as 1A-8A (i.e., Con 1C-2C = SCZ_GRNet modules with lower preservation in SCZ_nonGRNet and reverse, Con 3C-4C = Ht_GNet modules with lower preservation in Ht_nonGNet and reverse, Con 5C-6C = SCZ_GRNet modules with lower preservation in Ht_GNet and reverse, Con 7C-8C = SCZ_nonGRNet modules with lower preservation in Ht_nonGNet and reverse). ***C:*** Schematic representation of exploratory analyses performed to add biological plausibility to the co-expression “flexible” and “invariant” sub-networks: enrichments in gene ontologies (BP-biological processes, MF-molecular functions and CC-cellular components), PGC2-CLOZUK GWAS loci genes, height GWAS loci genes, SynGO and “druggable genome” genes.

### Generating variables for gene expression adjustment

This step included: a) estimation of variables used for data cleaning (quality surrogate variables or expression principal components); b) calculation of cell type proportions; c) computation of genomic scores for SCZ risk and height.

A prominent source of bias in re-constructing co-expression networks is RNA quality (i.e. technical or biological artifacts). To minimize the unwanted variance associated with this potential confounder, we implemented the recently described qSVA approach [22] (details in **SM2**). The resultant quality surrogate variables (qSVs) were subsequently used in the downstream analyses for adjusting the input for co-expression network analysis.

Cell type composition represents another factor that contributes to construction of co-expression networks from bulk RNA-Seq data, while modules of co-expression are likely to be significantly driven by cellular type [17]. Because in this study we sought to identify more subtle changes in gene correlatability associated with the genomic risk for schizophrenia, we opted to remove the variance explained by the relative cell type proportion. For this purpose, we used an approach implemented in the R package BRETIGEA (BRain cEll Type specIfic Gene Expression Analysis) [17, 23] (details in **SM3**). A resultant N_samples_ X M_cell proportion_ matrix, where the cellular types are represented by neurons (neu), astrocytes (ast), oligodendrocytes (oli), endothelial cells (end), microglia (mic) and precursors of oligodendrocytes (opc) was also used in the downstream analyses to adjust the input for co-expression network analysis.

c) Genomic scores from two GWAS studies-PGC2-CLOZUK [24] and height meta-analysis [25] - were calculated as previously described [5]. We used in this study the set of scores computed from SNPs that reached a significance level of p≤0.05 in each GWAS study. This is the 6^th^ set of standard scores based on GWAS p values, and although not representing the threshold of significance at GWAS level, it has the advantage of including a larger number of genetic variants potentially accounting for phenotypic variance in SCZ risk and height and generally assumes the asymptote of maximum risk accounted for by GRS. Accordingly, two scores were used in downstream analyses: Genomic Risk Score 6 (GRS6) for SCZ and Genomic Score 6 (GS6) for height (details in **SM4**).

### Variance analysis

We used functions from the *variancePartition* package [26] to assess the contribution of multiple variables to the expression variation of each gene (i.e., genomic risk score for schizophrenia (SCZ) and genomic score for height, age, sex, cell type proportion, qSVs (for LIBD data) or PCs (for CMC data), ethnicity (10 genomic principal components-snpPCs calculated from the genotype data), and technical parameters). Specifically, a multivariate linear (fixed effects) regression model was fit for each gene in the two data sets and the summary statistics were computed with the function *fitExtractVarPartModel*, including the variance fractions explained by each variable when controlling for all the other variables (**figure 1A**).

### Integration or depletion of genomic scores effects from the input for co-expression network construction

We used a data cleaning function-*cleaningY* available in the *jaffelab* package [27, 28] that allows removing the unwanted variance explained by variables of no interest and/ or technical confounders, while preserving the effect of variables of interest (**SM5**).

Residuals calculated from four models were used as expression input sets in weighted gene co-expression network analysis (WGCNA).

Accordingly, the four fitted models were:

1. Intercept and GRS6 schizophrenia “protected” (P=2 in cleaningY function);
2. Only intercept “protected” while GRS6 SCZ is removed with the other variables (P=1 in cleaningY function), i.e., GRS6 schizophrenia removed. Equation for the above 1-2 SCZ models:

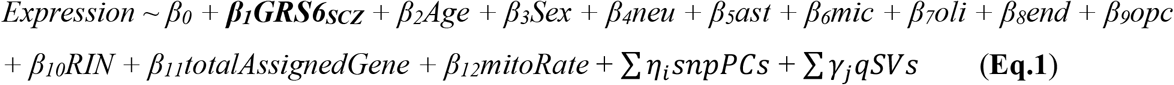
3. Intercept and GS6 height “protected” (parameter P=2 in cleaningY function);
4. Only intercept “protected” while GS6 height is removed with the other variables (P=1 in cleaningY function).

Analogous equations for the above 3-4 height models:

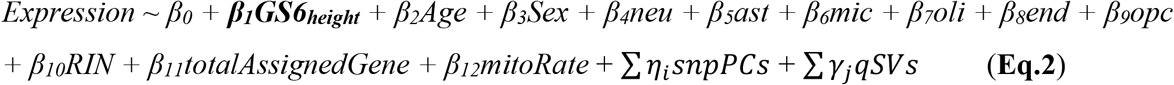

For both equations, i=10 snpPCs and j=8 qSVs.

Of note, for the CMC dataset, the first five principal components (calculated by *prcomp* function in R) for the expression data were used instead of qSVs in the above equations.

### Co-expression network analysis with WGCNA

The four new input datasets from each set of 78 LIBD and 116 CMC NT DLPFC tissue, representing the manipulation and variation of the effects of genomic risk for SCZ (as test) and genomic score for height (as negative control) on gene expression, were used for generating co-expression networks with WGCNA package [29] as previously described [5]. Briefly, gene pairwise correlations were computed (method = “bi-weight”), adjacency matrices were calculated (parameters: β power =6 estimated with the *sft* function, network type = “signed hybrid”), and modules of co-expressed genes were detected with hierarchical clustering (details about network construction in **SM6**).

By using the residuals from the models described above, two patterns of the co-expression networks were generated: 1. Networks of “preserved” genomic risk score (in statistical modelling jargon); in a more general sense networks “integrative” of susceptibility to modulatory influences by genomic signatures: “**SCZ G**enomic **R**isk co-expression **Net**work” (SCZ_GRNet) and “Height (**Ht**) **G**enomic profile co-expression **Net**work” (Ht_GNet); 2. Networks of “regressed out” polygenic risk score (again, in statistical modelling jargon); in a more general sense networks “depletive” of potential modulatory influences by the genomic signatures: “**SCZ non-G**enomic **R**isk co-expression **Net**work” (SCZ_nonGRNet) and “Height (**Ht**) **non-G**enomic profile co-expression **Net**work (Ht_nonGNet).

Of note, we further refer to the four networks as being either “integrative” or “depletive”, terms closer to their biological connotation.

We then interrogated the resultant co-expression networks with five aims:

I. Identification and general characterization of the “invariant” network structure (gene sets with the same gene content configuration in the “background” network irrespective of genomic modulatory effects).
II. Identification and characterization of areas in the “background” co-expression network that are “flexible,” i.e. show modulatory effects of polygenic risk.
III. Exploration of the biological plausibility of the network areas “flexible” or “invariant” with genomic signatures of SCZ risk and height within each dataset (intra-group validation).
IV. Evaluation of the networks’ relevance for SCZ pathophysiological mechanisms or drug targetable pathways.
V. Testing the consistency of “invariant” co-expression network architecture and areas “flexible” with genomic effects by comparisons between LIBD and CMC datasets (inter-group validation).

A general pipeline of the study design is presented in **figure 1**. We emphasize that, although we further refer to four networks per data set, they actually represent variant patterns of the same DLPFC gene co-expression network.

### I. Profiling the “invariant” background network

To isolate an “invariant” network architecture independent of the influence of the GRSs explored here, we used a cross-tabulation method implemented in the WGCNA *matchLabels* function. This function calculates the overlap between genes present in modules from a “source” and a “reference” network by computing a Fisher’s exact test. The modules of the “source” network are then re-labeled by the “reference” modules with which they have the most overlap. The source modules without a significant overlap in the reference network are considered specific for the source network and are given distinct labels. We applied this function by taking each network as reference one at a time and the other three as “source” networks. Consequently, a pairwise gene overlap was performed across the modules of the four networks and matched/ non-matched modules were identified for each network relative to the others. After matching the modules from the four networks, we extracted an “invariant” structure of the “background” network putatively unaffected by the genomic modulatory effects of SCZ and height, defined as all sets of minimum twenty genes overlaps from the matched modules across the four networks. The “invariant network” was extracted separately within each dataset, LIBD and CMC.

### II. “Flexible” gene sets of the co-expression network

To identify network areas that are “flexible” with the modulatory effects of genomic signatures of SCZ risk and of height, we planned “between traits” (BT) and “within traits” (WT) comparisons across the four networks for each dataset (LIBD and CMC) by using the gene overlap calculated with the *matchLabels* function and one network based statistics-connectivity - calculated with the WGCNA *modulePreservation* function. The planned comparisons are schematically represented in **figure 1B**.

Importantly, the contrasts used for differential network analysis should be regarded as context-dependent in which one network plays a dual and alternate role as “baseline” and “test” in reference to another network. Therefore, the outcome of a contrast is not necessarily network specific and rather singles-out patterns interpretable only in the context of how both networks are used for a comparison.

By pairwise gene overlap, we identified as “between traits” differential modules those modules that were unmatched modules from the contrasts SCZ_GRNet vs. H_GNet and reverse, which represent network differential configurations based on these specific genetic risk differences. The unmatched modules from the contrasts SCZ_nonGRNet vs. H_nonGNet and reverse represented network differential configurations based on residual variance of expression unexplained by the genomic signatures of traits of interest (SCZ or Ht) and nuisance covariates.

From the “within traits” contrasts, unmatched modules of “integrative” vs. “depletive” networks, respectively SCZ_GRNet vs. SCZ_nonGRNet and Ht_GNet vs. Ht_nonGNet contrasts were indicative of areas of network architecture vulnerable to modulatory genomic effects specifically of SCZ risk or height. Conversely, unmatched modules from SCZ_nonGRNet vs. SCZ_GRNet, respectively H_nonGNet vs. H_GNet contrasts were considered network architecture areas unexplained by modulatory effects from SCZ genomic risk or height genomic profile.

From an inferential perspective, the results of “between traits” contrasts represent patterns that show how far the modulatory effects of SCZ genomic risk on co-expression network deviate from those of a normative trait, i.e., height. Likewise, the outcome from contrasting SCZ_GRNet and SCZ_nonGRNet, likely carry the most relevant information about the effects of schizophrenia genomic risk on co-expression networks, i.e., illness risk element.

While calculating the gene overlap between network modules represents a simple and intuitive differential network analysis, other network-based statistics provide complementary and perhaps more nuanced qualitative differences between networks. Therefore, we also calculated the preservation statistics for the planned BT and WT contrasts by using the WGCNA *modulePreservation* function [30] (details in **SM7**).

From the module preservation connectivity output, we selected the median rank statistics recommended for modules with a wider range of sizes, while it does not depend on module size [30]. We selected connectivity, while it evaluates metrics related to degree, a well-known network parameter [30], showing to what extent the patterns of between-nodes connections are similar in a test versus a reference network [30-31]. Specifically, a higher value of median rank connectivity is indicative of lower similarity between the test and reference network [30]. Reference and test networks were designated in accordance with the planned comparisons (**figure 1**), respectively, we tested for the preservation of “integrative” networks’ modules in the “depletive” networks and vice-versa. Based on these contrasts, we selected as modules with evidence of lower preservation in the test network those within the 95^th^ percentile of the median rank connectivity scores. Of note, median rank preservation statistics generate relative measures, and therefore our cut-off should be regarded as exploratory and arbitrary. Moreover, while connectivity is rather a qualitative measure of similarity, it is possible for modules with high median rank connectivity to be part of the “invariant” co-expression sets, which are defined by different metric-the degree of gene overlap. Consequently, the modules selected based on preservation statistics were added to the unmatched modules from the cross-tabulation step only if they were not contributing to the “invariant network”.

In summary, the differential network analysis by cross-tabulation and module preservation statistics generated a total of 16 gene sets (unmatched modules and the less preserved modules based on the planned BT and WT comparisons), representatives of co-expression network areas variable with the presence or absence of modulatory effects from the genomic signatures of SCZ risk and height (**figure 1**). The 16 contrasts are henceforth referred as “flexible sub-network” sets. Each of the 16 contrasts includes the gene members of the differential modules for the corresponding contrast and represents an individual “flexible” sub-network set.

### III. Assessing the biological plausibility of the “invariant” and “flexible” sub-network sets

To identify potentially relevant biological mechanisms related to “invariant” and “flexible” sub-networks, we performed functional enrichment analysis with the g:GOSt function implemented in the gProfiler2 R package [32] (**SM8**). To highlight more specific “trait” related mechanisms, we also performed systematic comparative evaluations of the gene ontologies enriched in the 16 “flexible” sub-network sets. To this end, we used metrics of Gene Ontology Semantic Similarity, which allow comparisons of GO terms and GO annotated gene products [33]. Specifically, we compared the 16 “flexible” sub-networks in a pairwise manner by using the “Best Match Average” (BMA), a composite measure of similarity between GO lists, calculated with the “Wang” algorithm that leverages the Gene Ontology structure of Directed Acyclic Graph (DAG). For this analysis we used functions implemented in the GOSemSim R package [34] (details in the **SM9**). Briefly, we calculated the BMA between each “flexible” sub-network and all others and generated matrices of semantic similarity subsequently clustered and visualized through heatmaps created with the pheatmap R package [35].

### IV. Mapping gene sets of interest to “invariant” and “flexible” sub-networks

In addition to the exploratory research of biological plausibility associated with the “invariant” network architecture and “flexible” sub-network sets, we tested for enrichments in genes putatively more related to the two complex traits, respectively: PGC2-CLOZUK GWAS significant loci genes [24] (in short PGC2 genes), height GWAS significant loci genes [25] (in short “height” genes), and synaptic genes ontologies-in short SynGO genes [36]. Further, we aimed to map potentially druggable pathways to the networks by testing for enrichments in genes of the “druggable genome” [37]. Therefore, we used permutations to test hypotheses that “invariant” and “flexible” sub-networks are significantly enriched in several of the above genes of interest (details about permutations tests in **SM10**).

### V. Consistency and functional convergence of “invariant” and “flexible” sub-networks across datasets

For the inter-RNA-Seq dataset validation step we evaluated:

a) The overlap between LIBD and CMC “invariant” co-expression networks and between LIBD and CMC 16 “flexible” sub-networks, by using again permutation tests (N=10,000 iterations) (**SM10**).

b) The GO:BP semantic similarity between LIBD and CMC “invariant” co-expression networks and between LIBD and CMC 16 “flexible” sub-networks, by measuring the BMA as explained in section III and **SM9**.

The general assumption was that a significant overlap between pairs of corresponding LIBD and CMC “invariant”/ “flexible” sets, and/ or higher GO semantic similarity for the same comparisons, indicate consistency and functional convergence of the genomic profiles modulatory effects on the co-expression network architecture.

## RESULTS

LIBD and CMC samples’ demographics and expression data characteristics are represented in **figure 1**.

### Variance partition analysis

(**supplementary table 1, supplementary figures 1**-**2**) showed that in both data sets, GRS6 and GS6 accounted for minimal proportions of expression variance across the whole set of genes (**LIBD**-median variance by GRS6 SCZ = 0.59%; max = 26.6%; median variance by GS6 Ht = 0.32%, max = 12.6%; **CMC**-median variance by GRS6 SCZ = 0.45%, max = 11.4%, median variance by GS6 Ht = 0.46%, max = 13.4%); however, the explained variance was not uniformly distributed, with genes that significantly deviated from the median values for both scores. Moreover, the increasing variance explained by GRS6 or GS6 implicated different genes as suggested by the decreasing number of genes with higher variance explained by both genomic scores (**supplementary figure 2**).

### WGCNA results

The co-expression networks and modules based on expression inputs generated by varying the integration of GRS6 SCZ and GS6 height are summarized in **figure 1** and represented in the cluster dendrogram-heatmap plot from **supplementary figure 3**.

### The “invariant” and “flexible” sub-networks profiles

The output of cross-tabulation showed that 24 LIBD clusters (Total =2596 genes; set size: 23-380) and 38 CMC clusters (Total = 3344 genes; set size 20-375) qualified as “invariant”, potentially unaffected by modulatory effects of genomic profiles for SCZ risk and height (see **supplementary table 2** for the gene membership in the “invariant” sets). Overall, the LIBD and CMC “invariant” sub-networks shared 582 genes representing a statistically significant overlap by permutation tests (**supplementary figure 4**).

Planned comparisons generated 16 gene sets considered as areas of the co-expression network putatively influenced by SCZ genomic risk or height genomic profile (“flexible” network). The LIBD flexible network associated contrasts (size = 104-768 genes) and CMC contrasts (size = 99-1894) included a total of 3634 (LIBD), and 5211 (CMC) unique genes, respectively (the full gene composition of contrasts is presented in **supplementary table 3**). A total of 1116 genes were members of “flexible” networks in both datasets (LIBD and CMC) and the overlap was significant by permutations test (p=0.0004). Of note, the contrasts were not perfectly distinct with roughly 54% of genes in LIBD contrasts (N=1946) and approximately 28% of genes in CMC contrasts (N=1467) being members of more than one contrast. Collectively, these observations suggest again that both “invariant” and “flexible” sub-networks are context-dependent and relative; consequently, their meaning is uncertain without evidence of biological plausibility and additional empirical data.

### Biological plausibility of the “invariant” sub-network

The enrichment of “invariant” gene co-expression areas, i.e. areas not influenced by genetic risk for schizophrenia or for height, in meaningful ontologies suggested that they belong to functionally coordinated and critical pathways, which recapitulate biological motifs reported in earlier work [5, 10, 38], methodological differences notwithstanding (i.e., the control for cell type proportion). In both datasets, “invariant” network sets were enriched for a) biological processes specific to the nervous system development and functionality (e.g., nervous system development, modulation of chemical transmission and signaling) and b) fundamental, tissue unspecific processes (e.g., transcription, translation and metabolism) (**supplementary table 4**). Moreover, permutations results showed that 15 LIBD and 20 CMC “invariant” clusters had significant overlap (**figure 2A, supplementary table 5**). Further, the GO biological processes associated with all but six overlapped clusters (4 LIBD and 2 CMC) indicated varying degrees of GO semantic similarity ranging from weak to strong (**figure 2B**). By GO semantic similarity, the “invariant” overlapping LIBD and CMC clusters aligned along three functional dimensions represented by pathways specific to nervous system intertwined with more general cellular processes (**figure 2C, D, E**).

**Figure 2:**
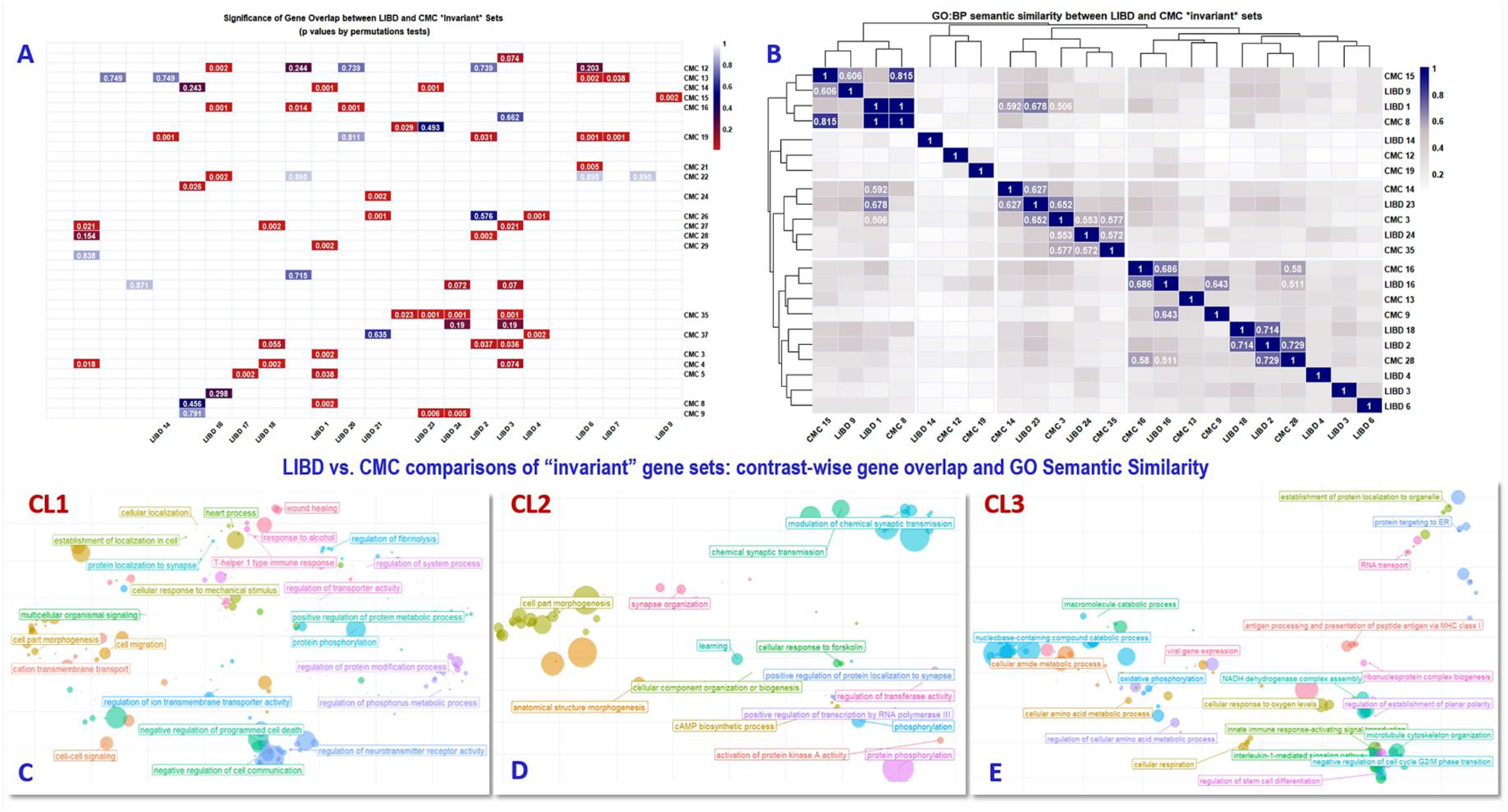
LIBD vs. CMC comparisons of “invariant” gene sets: contrast-wise gene overlap and GO semantic similarity. ***A:*** Heatmap representing LIBD-CMC corresponding “invariant” sets. ***B:*** Significantly overlapped “invariant” sub-networks were also tested for GO biological processes semantic similarity by comparing GO:BP terms enriched in LIBD and CMC “invariant” sets; cluster heatmap shows a relative hierarchization of corresponding LIBD-CMC “invariant” sub-networks by GO:BP semantic similarity measured with BMA, respectively LIBD-CMC “invariant” sets with BMA ≥0.5 are organized in three clusters; GO:BP terms enriched in the three clusters are represented in panels C-E. Visualization of heatmap plots in panels A-B was created with R package pheatmap [35] and GO:BP scatter plots in panels C-E were created with R package rrvgo [48].

Interestingly, testing enrichments of “invariant” networks in gene sets of interest showed that no “invariant” cluster was significantly enriched in PGC2 CLOZUK genes. There were, however, various “invariant” clusters enriched in GWAS height genes, SynGO and druggable genome genes (**supplementary table 6**).

### Biological plausibility of the “flexible” sub-networks

All 16 contrasts in LIBD and CMC datasets, i.e., those contrasts illuminating network features that are modulated by genetic risk for either schizophrenia or height and deemed as “flexible” sub-networks, were enriched for gene ontologies of reasonable interest. Interestingly, by GO:BP (biological processes), only the “flexible” sub-networks having SCZ_GRNet as one term of comparison, respectively three LIBD (WT contrasts 1C, 2A and BT 5A) and one CMC “flexible” sub-networks (BT 6A) were enriched for CNS ontologies previously highlighted in genetic and molecular studies of schizophrenia (i.e., synaptic signaling and ion transport, neuron development and axon genesis, etc.) [39]. In both RNA-Seq datasets, “within traits” contrasts related to height, i.e. GS6_Ht vs. GS6_nonHt networks, were enriched for general cellular processes mainly related to mRNA metabolic processes and protein translation (see **supplementary table 7** for complete gene ontology enrichment), GO terms not generally associated with schizophrenia.

Perhaps surprisingly, none of the LIBD or CMC contrasts based on flexible areas were significantly enriched for PGC2 CLOZUK GWAS loci genes (**figure 3**). Further, only two WT contrasts were significantly enriched for meaningful gene sets of interest across the two datasets: Con2A (SCZ_nonGRNet vs. SCZ_GRNet) was enriched for SynGO genes and Con 3A (Ht_GNet and Ht_nonGNet) was enriched for height GWAS genes (**figure 3**).

**Figure 3:**
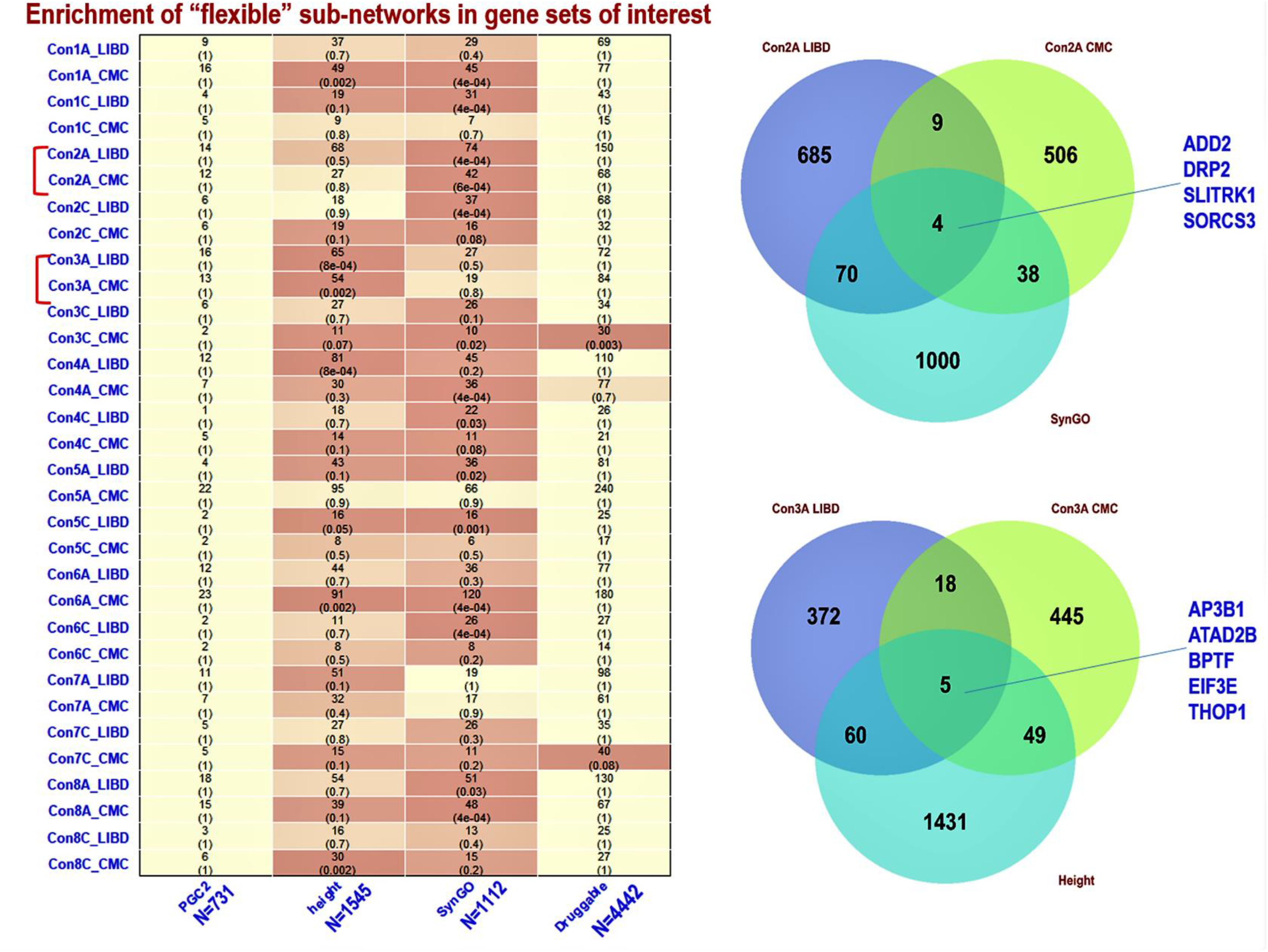
Mapping gene sets of interest to “flexible” sub-networks. Only two contrasts were significantly enriched for relevant gene sets in both LIBD and CMC datasets: Con2A (within traits: SCZ_nonGRNet vs. SCZ_GRNet) enriched in genes from synaptic ontologies and Con3A (within traits Ht_GNet vs. Ht_nonGNet) enriched in GWAS height loci genes. The Venn diagrams show the size of overlap and the genes shared by the two contrasts and the respective gene set of interest, i.e., SynGO and Con2A, height genes and Con3A.

**Figure 4:**
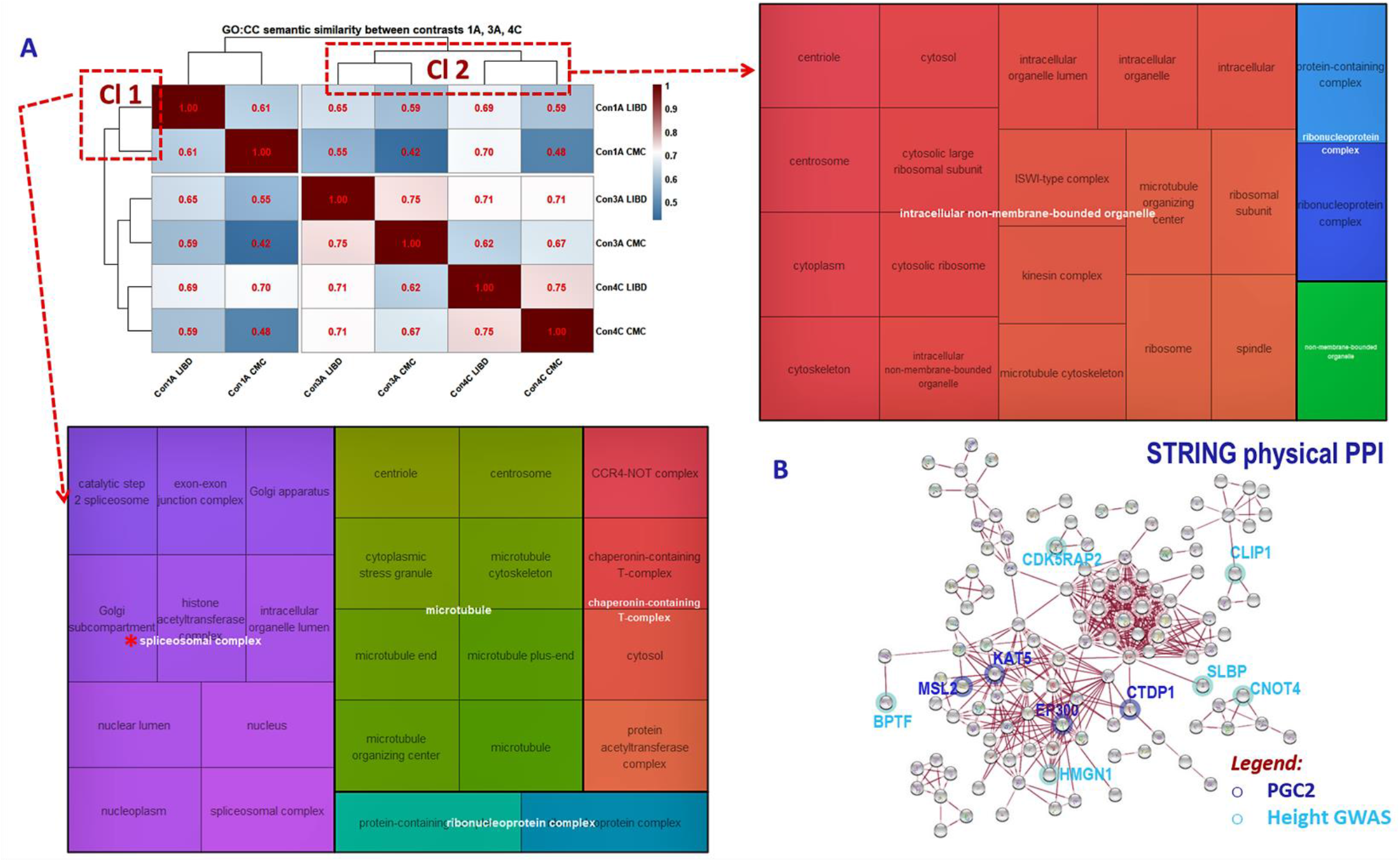
GO cellular component (GO:CC) semantic similarity between LIBD and CMC “flexible” sub-networks. ***A:*** SCZ and “height” based networks show a trend for segregation as suggested by the cluster heatmap of GO:CC semantic similarity (upper left panel). Upper right and bottom left panels: while the two clusters (Cl1 and Cl2) share several GO cellular components, one GO:CC term-spliceosome complex - differentiates between them. Visualization of heatmap plots was created with the R package pheatmap [35] and the treemap plots with the R package rrvgo [47]. ***B:*** Genes from the GO:CC ontologies differential between Cl1 and Cl2, implicated in protein-protein interactions (PPI) networks (see **supplementary material** for details about gene filtering criteria and PPI network calculation with STRING).

To assess the potential consistency between LIBD and CMC “flexible” sub-networks, we used two criteria: significant gene overlap by permutations tests and global BMA ≥ 0.5 in at least one gene ontology domain. Based on these criteria, we identified three WT and one BT contrast: Con 1A, 3A and 4C, respectively Con 8C.

Overall, the three WT contrasts (1A, 3A and 4C) showed an apparently weaker degree of consistency between LIBD and CMC (**table 1**). However, when comparing the GO:CC between them and across datasets, in spite of the seemingly negligible differences in BMA, Con 1A LIBD and CMC clustered together, while Con 3A and 4C (both WT contrasts based on Ht_GS6 and Ht_nonGS6) formed a separate cluster (**figure 3A** left top panel). To further understand what cellular component distinguishes between Con 1A and Con 3A, 4C across datasets, we examined the GO:CC with higher similarity (BMA ≥ 0.5) in each cluster (1A LIBD - 1A CMC, respectively 3A, 4C LIBD - 3A, 4C CMC) and we found that GO:CC “spliceosomal complex” is specific to the 1A LIBD - 1A CMC cluster (**figure 3A** bottom left panel). We then isolated the genes associated with GO:CC in 1A LIBD - 1A CMC cluster and examined them in STRING PPI database to identify participants in protein-protein physical interactions as indicated by experimental evidence [40] (**figure 3B**). Notably, four of the PPI members are within PGC2 GWAS loci genes (CTDP1, EP300, MSL2 and KAT5) and six PPI members are within the GWAS height loci (CLIP1, SLBP, CNOT4, HMGN1, BPTF and CDK5RAP2), underscoring the interactive contribution of genomic risk for SCZ and the genomic profile for height to the brain co-expression and subsequently protein-protein interaction networks.

**Table 1:** Gene overlap (permutations tests) and Gene Ontology Semantic Similarity between LIBD and CMC “flexible” sub-networks.

| Gene overlap |  |  | GO Semantic Similarity (BMA) |  |  |
| --- | --- | --- | --- | --- | --- |
| Contrasts | Set size (LIBD/CMC) | Overlaps (p values) | GO:BP | GO:MF | GO:CC |
| Con 1A | 463/541 | 17 (0.017) | 0.195 | 0.097 | 0.607 |
| Con 1C | 158/183 | 0 (1) | NA | 0.203 | 0.362 |
| Con 2A | 768/557 | 13 (0.74) | 0.388 | 0.557 | 0.578 |
| Con 2C | 312/244 | 5 (0.18) | 0.819 | 0.399 | 0.762 |
| Con 3A | 455/517 | 23 (0.004) | NA | NA | 0.749 |
| Con 3C | 319/99 | 1 (0.74) | 0.162 | 0.366 | 0.431 |
| Con 4A | 633/448 | 16 (0.06) | 0.128 | 0.2 | 0.529 |
| Con 4C | 229/150 | 15 (0.004) | 0.39 | 0.129 | 0.746 |
| Con 5A | 404/1894 | 24 (0.74) | 0.354 | 0.411 | 0.575 |
| Con 5C | 104/124 | 1 (0.47) | NA | 0.143 | 0.506 |
| Con 6A | 543/1162 | 45 (0.004) | 0.083 | 0.193 | 0.315 |
| Con 6C | 129/120 | 2 (0.173) | 0.291 | 0.179 | 0.543 |
| Con 7A | 497/524 | 16 (0.04) | NA | 0.081 | 0.303 |
| Con 7C | 370/183 | 4 (0.28) | 0.637 | 0.576 | 0.694 |
| Con 8A | 638/545 | 25 (0.02) | 0.312 | 0.632 | 0.246 |
| Con 8C | 198/275 | 10 (0.004) | 0.068 | NA | 0.866 |

In summary, the results of our novel approach in constructing brain co-expression networks by varying the expression input with genomic profiles of schizophrenia risk and height generated several key observations: 1) Non-negligible variability of co-expression modules with integration or removal of residual variance explained by GRS6 SCZ and GS6 height; 2) Biological plausibility and potential relevance for illness etiopathogenesis of gene sets isolated by comparing co-expression networks; 3) The lack of clustering of GWAS loci genes in specific modules or sub-networks and 4) The sparse replicability of findings in the two independent datasets.

## DISCUSSION

The purpose of this study was to explore how integration of particular genetic contexts in gene expression inputs would modulate the co-expression network architecture in the neurotypical brain. Specifically, we tested this approach to gain deeper and possibly novel perspectives about the likely modulatory effects of polygenic risk for schizophrenia on co-expression networks in contrast with a normative control trait, i.e., height. If the co-expression network from the postmortem brain is conceptualized as a combination of dynamic signaling interactions, it is conceivable that its construction will capture specific aspects depending on modelling various contexts including genomic profiles for pathological or normative traits. Furthermore, contrasting networks with different genetic signatures would isolate network characteristics relevant for various traits. However, we emphasize the particular and qualitative nature of such comparisons, dependent on the entire context of expression input manipulation and consequently the difficulty of generalizations. Therefore, we regard this approach as exploratory for uncovering snapshots of context-dependent network configurations whose findings need to be interpreted through the angle of biological plausibility and validated by further experimental approaches.

In light of this perspective, the main findings of our study are as follows: 1) a consistent pattern of gene co-expression in DLPFC of neurotypical brain unmodulated by genetic risk for schizophrenia or height (i.e., “invariant” gene sets) combined with subtle conformational variability owing to slight changes in the expression input; 2) the presence of network areas susceptible to and “flexible” with genomic variation associated with schizophrenia or height; 3) notwithstanding the expression input manipulation, the “invariant” and “flexible” sub-networks displayed biological plausibility as suggested by gene ontology analysis and/ or enrichments in gene sets of interest; 4) organization of biological ontologies associated with “flexible” sub-networks/ contrasts along contiguous dimensions varying between low GO semantic similarity (which tag “distinct” contrasts) and high GO semantic similarity (related to more “fuzzy” contrasts); 5) a degree of gene overlap between “flexible” sub-networks (contrasts) modulated by schizophrenia risk and height genomic profiles, an observation consistent with their relative nature, and possibly explained by the pleiotropic character of genes that can affect multiple complex traits by overlapping mechanisms; 6) in spite of differences in methodology and gene expression characteristics, several results demonstrated various levels of consistency in an independent dataset, though consistency is notably limited. Several “invariant” network sub-sets consistent across LIBD and CMC datasets demonstrated gene overlap more significant than expected by chance, but most importantly showed biological plausibility by three clusters of Gene Ontology semantic similarity that captured fundamental biological processes in combination with specialized CNS functions.

A similar but less consistent pattern of mixed GO:BP also characterized the “flexible” sub-networks. This observation is potentially important for two reasons: 1) it suggests that at least fragments of biological pathways are “modulated” by polygenic profiles of complex traits; 2) the potentially affected pathways are not circumscribed to one mechanism, nor do they segregate, but are rather intermixed and show “fuzzy” boundaries. The admixture of cellular biological processes associated with the “flexible” sub-networks underscore the plausible “omnigenic” profile of schizophrenia risk [41], embedded within the mesh of multiple other complex traits. Notwithstanding the pattern of mixed biological processes, the more CNS related GO:BP were rather enriched in LIBD/ CMC contrasts derived from the GRS6 “integrative” network (SCZ_GRNet) as term of comparison.

As mentioned above, the within trait (WT) contrasts/ “flexible” sub-networks that capture the modulatory effects of GRS6 SCZ are potentially the most informative from a biological perspective related to schizophrenia. While at first blush the biological processes consistency across our two datasets looks disappointing, at closer inspection Con 1A shows relative specificity toward the “spliceosomal complex”, a cellular component plausibly disrupted in schizophrenia and other neuropsychiatric disorders, as highlighted in previous work from our group and others (12, 42-46). Whereas the transcriptional regulation through alternative splicing is a pervasive mechanism affecting hundreds of genes on different pathways, perhaps underwriting some of the heterogeneity of schizophrenia, it should come as no surprise that the “flexible” sub-networks “integrative” of genomic risk for SCZ show less consistency in terms of biological processes across the two different datasets that we explored. The LIBD and CMC brain RNAseq data come from brains significantly different by mean age and from analyses with differing approaches to mRNA quality correction and other QC procedures.

Of interest from the perspective of SCZ genomic risk and potentially drug target development, four genes in Con 1A involved in transcriptional regulation and implicated in protein-protein physical interactions, are within the PGC2 GWAS loci: KAT5 (Lysine Acetyltransferase 5), EP300 (E1A Binding Protein P300), MSL2 (MSL Complex subunit 2) and CTDP1 (CTD Phosphatase subunit 1). Moreover, these genes are implicated in various neurodevelopmental pathways or diseases [47]. We find it particularly noteworthy, however, that although several PGC2 loci genes were mapped to “flexible” sub-networks of interest as illustrated in the example above, we did not see a trend of clustering of GWAS genes in particular sub-networks. While perhaps surprising, as these flexible networks were constructed based on GRS, this observation underscores that genomic risk of schizophrenia is probably not limited to GWAS hits and represents an extensive background on which complex gene-gene, gene-environment or pathways interactions concur in the schizophrenia development. Prior studies of gene co-expression networks and their relationship to GWAS genes were based on enrichment statistics and correlation with modules’ eigengenes [5, 14]. Our new approach proposed in this study is a potentially more granular view to understand how genetic risk influences the behavior of signaling pathways in cells because we created the networks based on the influence of genetic risk.

Notwithstanding the broad co-expression network characterization, our study is not without limitations. One important caveat is related to the generalizability of results. Although we used comprehensive regression models to control for multivariate effects on gene expression with further consequences on co-expression networks and independent datasets, by design unaccounted factors could explain better than genomic profiles the networks configurations. For example, a possible limitation is related to the selection of the genomic profile of height as a “negative control” for comparison with genomic risk for SCZ in terms of effects on co-expression network architecture. While SCZ and height are seemingly two unrelated traits, they most probably share some common biological pathways.

We also used rather arbitrary, perhaps too lenient thresholds both for the selection of genomic scores (GRS6 SCZ and GS6 height) and for filtering the modules with “low preservation”. However, we chose these selections based on our interest to capture a larger proportion of variability and by the heuristic nature of our study. Also of note, the filtering procedure for connectivity preservation, combined with the not negligible number of grey genes resulted from the WGCNA, left a lower number of nodes/ genes as components of the network architecture. However, the strength of our study doesn’t come merely from proving the variability of brain co-expression network with biological and technical factors, but it also suggests that novel, well designed network analysis methods will be necessary to better capture the complexity of the conceivable dynamic co-expression networks in the brain.

Another potential limitation is the sample size of the postmortem brain data we used in our co-expression network re-construction in comparison with studies based on consortium data (i.e., PsychENCODE) [9, 12]). However, we used more homogeneous sample including only subjects with European ancestry in order to avoid population stratification confounds when creating the co-expression networks; likewise, because of more consistent tissue processing procedure, we believe our findings are more robust.

In conclusion, our study provides a novel characterization of postmortem, “neurotypical” DLPFC gene co-expression networks, part of which seems plausibly modulated by genomic signatures of normal or pathological complex traits. Further studies at various levels, conceptual, methodological (novel network analysis techniques), and especially experimental, are warranted for disentangling genomic and environmental interactions contributing to the risk of schizophrenia.

## Acknowledgements

We are grateful for the generosity of the Lieber and Maltz families in establishing an institute dedicated to understanding the basis of developmental brain disorders. We would like to gratefully acknowledge the families of the subjects whose donations made this research possible. This research was funded by the Lieber Institute for Brain Development.

Dr. Eugenia Radulescu is grateful to Dr. Pasquale Di Carlo, Dr. Andrew Jaffe and Dr. Richard Straub for analytical advice and invaluable comments on earlier versions of this manuscript.

## FIGURES AND TABLES LEGENDS

**Supplementary table 1:**Gene-wise distribution of variance explained by the regressors from models used to adjust the expression input for WGCNA.

**Supplementary table 2:**Sub-networks “invariant” with genomic profiles of SCZ and height, derived from LIBD and CMC co-expression networks.

**Supplementary table 3:**Sub-networks “flexible” with modulatory effects of genomic profiles of SCZ and height, isolated from LIBD and CMC co-expression networks.

**Supplementary table 4:**“Invariant” sub-networks enrichment in Gene Ontology terms and annotations performed with gprofiler.

**Supplementary table 5:**Consistency between LIBD and CMC “invariant” sub-networks by gene overlap tested with permutations tests.

**Supplementary table 6:**“Flexible” sub-networks enrichment in Gene Ontology terms and annotations performed with gprofiler.

**Supplementary table 7:**“Flexible” sub-networks enrichment in PGC2-CLOZUK GWAS loci, “height” GWAS loci, SynGO and “druggable genome” genes.

**Supplementary figure 1:**Characterization of gene expression variance profiles converging from multifactorial effects on DLPFC transcriptome within the two data sets/ groups (LIBD, CMC).

**Supplementary figure 2:**QQ plots and Venn diagrams showing the gene-wise divergence between the increasing variance explained by GRS6 SCZ and GS6 height.

**Supplementary figure 3:**Matched modules across four co-expression networks calculated with WGCNA based on expression inputs that “integrate” or “deplete” effects of GRS6 SCZ and GS6 height.

**Supplementary figure 4:**The null distribution for the overlap of two gene sets with the size of “invariant” LIBD and CMC sub-networks (hypergeometric and 10,000 permutations tests).

**Supplementary figure 5:**Venn diagram showing the total HGCN annotated genes overlap between LIBD and CMC data sets.

