## Supplemental material for "Investigating trait variability of gene co-expression network architecture in brain by manipulating genomic signatures of schizophrenia risk"

**Supplementary material**

**Supplementary methods (SM)**

**SM1: Postmortem brain samples and RNA-Seq processing**

*LIBD RNA-Seq*

Postmortem brain tissue was collected, dissected and processed under a protocol described in Jaffe et al [1], and Collado-Torres et al [2].

RNA sequencing: total RNA was extracted from DLPFC gray matter (BA9/46) with RNeasy Lipid Tissue Mini Kit (QIAGEN) and sequencing libraries were constructed with the TruSeq Stranded Total RNA Library Preparation kit with Ribo-Zero Gold ribosomal RNA depletion.

Raw sequencing reads were quality checked with FastQC [3] and corrected with Trimmomatic [4] if necessary. The quality checked sequencing reads were mapped to the hg38/GRCh38 human reference genome with HISAT2 (v2.0.4) [5]; following alignment, the expression for genes and exons was summarized in counts based on GENCODE v25 (GRCh38.p7) [6] and converted to RPKM (Reads Per Kilobase of transcript per Million mapped reads).

Genes with sufficient abundance (RPKM ≥0.1) in more than 80% of samples (N=18,980 genes) were then normalized by log_2_(x+1) transformation. Lastly, we removed the samples outlying for the standardized connectivity, which was computed by hierarchical clustering of the Euclidean distances measured from the expression data [7].

*CMC RNA-Seq*

CMC expression data release 1 was downloaded from Synapse (<https://www.synapse.org/>) and processed as previously described [8] and the FASTQ files were used in the SPAEQeasy pipeline [9], which consists of the following steps: alignment to the hg38/GRCh38 human reference genome with HISAT2 (v2.0.4), pseudo-alignment to a reference transcriptome with kallisto or salmon [10-11], expression features (i.e., genes) quantified with featureCounts [12] and regtools [13], assembling the SummarizedExperiment R objects (the output of SPEAQeasy).

Then, similarly with LIBD expression data, genes with sufficient abundance (RPKM ≥0.1) in more than 80% of samples (N=27,805 genes) were normalized by log_2_(x+1) transformation. Finally, we removed the samples outlying for the standardized connectivity, which was computed by hierarchical clustering of the Euclidean distances measured from the expression data [7].

**SM2: Calculation of quality surrogate variables for removing unwanted variance from RNA quality**

Quality Surrogate Variable Analysis (qSVA) consists of identifying the transcript features most susceptible to RNA degradation, respectively, estimating the so-called “degradation matrix” quantified as the coverage of the susceptible features in customized sequencing libraries and performing a principal component analysis on the “degradation matrix” that yields a number of k principal components named “quality surrogate variables” (qSVs).

Quality surrogate variables (qSVs) were computed from a RNA degradation matrix as described in Jaffe et al [14].

The degradation matrix consisted of expression measures for 1000 chromosomal regions most susceptible to RNA decay when brain tissue from 5 donors was exposed to the room temperature for various intervals of time [14]). Quality-surrogate variables computed from the degradation-matrix (“degradation principal components”- qSVs hence fort) were derived with the sva package in R, which implements a principal component analysis algorithm.

The number of qSVs was pre-determined with the num.sv function that uses the method of Buja and Eyuboglu (option method=”be” in the num.sv() function).

**SM3: Calculation of cell type proportion from bulk RNA-Seq data with BRETIGEA**

**BR**ain c**E**ll **T**ype spec**I**fic **G**ene **E**xpression **A**nalysis implements a singular value decomposition algorithm to calculate a sample specific cell type proportion based on genes previously determined as cell type markers by meta-analysis [15].

The program takes bulk RNA sequencing data from brain regions (in our case DLPFC) and performs a brain cell type proportion estimation analysis that generates a matrix of surrogate proportion variables corresponding to six major brain types (neurons, astrocytes, oligodendrocytes, microglia, endothelial cells and oligodendrocytes precursors). A specific parameter allows the user to select the number of marker genes. For our study we used the default nMarker=50 (see details about trade-off when selecting the number of markers in the BRETIGEA user manual [16]). The matrix of cell type proportions was subsequently used in our models to adjust the gene expression input for WGCNA.

**SM4: GRS calculation**

GRS for schizophrenia were calculated for each individual as a measure of genomic risk for schizophrenia, based on the most recent published schizophrenia GWAS study [17]. We obtained odds ratios of 103,563 index SNPs from a meta-analysis of CLOZUK+PGC2 GWAS. These 104K SNPs are LD independent (R^2^<0.1) and span across the whole genome. We then calculated a weighted sum of risk alleles for schizophrenia, by summing the imputation probability for the reference allele of the index SNP, weighted by the natural log of the odds ratio of association with schizophrenia, at each independent locus across the whole genome, as described elsewhere [18, 19]. Consistent with the original approach taken by the Psychiatric Genomics Consortium in the GWAS study [19], ten PRS (PRS1-PRS10) were calculated using sub-sets of the 104K SNPs under different thresholds of the PGC2 GWAS p-values of association with schizophrenia: 5e-08, 1e-06, 1e-04, 0.001, 0.01, 0.05, 0.1, 0.2, 0.5, and 1.

**SM5: Data cleaning with cleaningY() function**

The *cleaningY* function available in the jaffelab R package (version 0.99.30) (https://github.com/LieberInstitute/jaffelab) fits linear regression models with expression data as dependent variables and multiple variables as independent regressors and removes the variance associated with variables of non-interest (for the current study age, sex, cell-type proportion and ethnicity represented by 10 genomic PCs), hidden (qSVs or PCs) and observed technical confounders (RIN, mithocondrial mapping rate and total assigned gene rate), while keeping (“protecting”) the intercept and variables of interest (in this study, genomic risk score for SCZ or genomic score for height). In other words, “protected” effects are estimated but not marginalized.

**SM6: WGCNA**

For the co-expression network analysis we used WGCNA package version 1.68 [20]. Adjusted expression data were used as input for network construction. Before selecting the beta power, the outliers for expression were removed as specified in **SM1**.

The beta power threshold was selected by analysis of scale free topology for multiple soft thresholding powers (sft function in WGCNA).

The networks were automatically created with *blockwiseModules* function. Parameters: correlation type = bi-weight midcorrelation; type of network = signed hybrid; power=6 selected with soft thresholding to correspond to an R^2≥0.8; merging the modules whose eigengenes are highly correlated for a default height of 0.15; minimum module size=20, PAM stage = TRUE, respectively, performing a second module detection PAM-like.

Modules were detected with a hierarchical clustering algorithm and labeled with pre-specified colors implemented in WGCNA routines.

**SM7: WGCNA module preservation**

Network based preservation statistics calculated with modulePreservation function in WGCNA is summarized in composite scores and median ranks of three network’s characteristics: density, connectivity and separability. This function measures how well the modules of a reference set are preserved in a test set.

For our study we specifically selected connectivity, while it is derived from metrics related to a well-known network parameter, “degree”, defined as the sum of gene’s connections strengths with other genes (a.k.a. “nodes”) in the network (tables 1 and 2 in Langfelder et al [21]).

We also selected the median rank connectivity that is better suited for modules with different sizes, as recommended in Langfelder et al [21], whereas it is based on observed values not on Z summary scores.

For this study we used modulePreservation function with parameters: network type = “signed hybrid”, corFun = “bicor”, nPermutations=1.

The reason for using just one permutation is that median rank statistics are based on observed values and a p value is not calculated, consequently it is recommended to turn-off the number of permutations to avoid the computational burden associated with permutations tests [21].

**SM8: Gene Ontology analysis**

Functional enrichment of modules or gene sets of interest was performed with the R version of gProfiler, respectively gprofiler2 v.0.2.0. Specifically, the g:GOSt function implemented in gprofiler2 was used.

This function performs a hypergeometric test by which functional terms (i.e., gene ontology sources such as biological process- GO-BP, molecular function- GO-MF, cellular components- GO-CC) are tested for statistically significant enrichment in gene lists of interest [22]. For multiple comparisons correction g:GOSt uses a custom algorithm- gSCS (Set Counts and Sizes) that accounts for the overlap of functional terms [22]. Likewise, we employed the g:GOSt function by using as statistical domain a custom background represented by: the overlap between LIBD, CMC and genes annotated by HGCN symbol downloaded from <https://biomart.genenames.org/>, the gene overlap between LIBD and CMC, the gene overlap between LIBD and HGCN genes and the gene overlap between CMC and HGCN sets (see the Venn diagram in supplementary figure 5). A total of 18,301 annotated genes were in the background used for functional enrichment analysis.

**SM9: GO Semantic Similarity Analysis**

To asses GO semantic similarity between gene sets of interest we used GOSemSim R package that implements methods for measuring semantic similarity between GO terms and gene products [23].

Specifically, we compared lists of GO terms enriched in gene sets of interest by using the “Wang method”, which leverages the Gene Ontology structure of DAG (Directed Acyclic Graph).

By this method, a GO term specificity is determined by its location on the GO graph. In other words, this algorithm is a graph-based strategy that measures semantic similarity using the topology of the GO-graph structure.

As a composite measure of similarity between the GO lists, BMA (**B**est **M**atch **A**verage) was selected; BMA represents the average of all maximum similarities on each row and column of a similarity matrix (see formula in Yu et al [23]). BMA takes values between 0-1 and it suggests small similarity for values of 0.5, medium similarity for values of 0.7 and large similarity for values around 0.9.

**SM10: Permutations tests**

To determine the significance of overlap between gene sets of interests we used permutations tests performed with functions implemented in R package “purrr”.

Specifically we used 10,000 permutations to generate the null distribution for the overlap between two gene sets with the same size as the original gene sets (i.e., individual “invariant” clusters, or “flexible” gene sets and PGC2/ “height”/ “SynGO”/ “druggable”) by randomly sampling genes without replacement from the full gene list (i.e., the group specific DLPFC set of genes plus the genes of interest not present in the DLPFC dataset), followed by computing the number of overlapping genes. To assess the statistical significance, we calculated the p values representing the number of simulated values greater than or equal to the observed values (plus one) divided by the number of iterations (plus one). Depending of the context in which permutations tests were used, we also corrected for multiple comparisons. For example, if testing for enrichment in gene sets of interest (PGC2, SynGO, etc.), due to the repeated testing of the respective gene sets (i.e. 16 times in the number of contrasts, or times number of “invariant” clusters/ per datasets), a multiple comparisons adjustment correction was used (p.adjust function, with FDR method).

### 8. Jaffe AE, Straub RE, Shin JH, Tao R, Gao Y, Torres LC, et al. Developmental and genetic regulation of the human cortex transcriptome illuminate schizophrenia pathogenesis. *Nat Neurosci*. 2018 August; 21(8):1117-1125.

9. Eagles et al: <https://www.biorxiv.org/content/10.1101/2020.12.11.386789v1>

10. Bray NL, Pimentel H, Melsted P, Pachter L. Near-optimal probabilistic RNA-seq quantification. Nat Biotechnol. 2016 May 01;34(5):525-7.

11. Patro R, Duggal G, Love MI, Irizarry RA, Kingsford C. Salmon provides fast and bias-aware quantification of transcript expression. Nat Methods. 2017 April 01;14(4):417-9.

12. Liao Y, Smyth GK, Shi W. featureCounts: an efficient general purpose program for assigning sequence reads to genomic features. Bioinformatics. 2014 April 01;30(7):923-30.

13. Y.-Y. Feng *et al.* , RegTools: Integrated analysis of genomic and transcriptomic data for

discovery of splicing variants in cancer. *BioRxiv* (2018), doi:10.1101/436634.

14. Jaffe AE, Tao R, Norris AL, Kealhofer M, Nellore A, Shin JH, et al. qSVA framework for RNA quality correction in differential expression analysis. *Proc Natl Acad Sci U S A*. 2017 July 03;114(27):7130-5.

15. McKenzie AT, Wang M, Hauberg ME, et al. Brain cell type specific gene expression and co-expression network architectures. *Sci Rep*. 2018;8(1):8868-5. doi: 10.1038/s41598-018-27293-5 [doi].

16. BRETIGEA: <https://cran.r-project.org/web/packages/BRETIGEA/index.html>

17. Pardinas AF, Holmans P, Pocklington AJ, et al. Common schizophrenia alleles are enriched in mutation-intolerant genes and in regions under strong background selection. *Nat Genet*. 2018;50(3):381-389. doi: 10.1038/s41588-018-0059-2 [doi].

18. International Schizophrenia C, Purcell SM, Wray NR, Stone JL, Visscher PM, O'Donovan MC*, et al.* Common polygenic variation contributes to risk of schizophrenia and bipolar disorder. Nature 2009; 460(7256): 748-52.

19. Schizophrenia Working Group of the Psychiatric Genomics C. Biological insights from 108 schizophrenia-associated genetic loci. Nature 2014; 511(7510): 421-7.

20. WGCNA: <https://cran.r-project.org/web/packages/WGCNA/WGCNA.pdf>

21. Langfelder P, Luo R, Oldham MC, Horvath S. Is my network module preserved and reproducible? *PLoS Comput Biol*. 2011 January 20;7(1):e1001057.

22. Raudvere U, Kolberg L, Kuzmin I, et al. G:Profiler: A web server for functional enrichment analysis and conversions of gene lists (2019 update). *Nucleic Acids Res*. 2019;47(W1):W191-W198. doi: 10.1093/nar/gkz369 [doi].

23. Yu G, Li F, Qin Y, Bo X, Wu Y, Wang S. GOSemSim: an R package for measuring semantic similarity among GO terms and gene products. Bioinformatics. 2010 April 01;26(7):976-8
