## Supplementary figures 1-5 for "Investigating trait variability of gene co-expression network architecture in brain by manipulating genomic signatures of schizophrenia risk"

### Slide 1
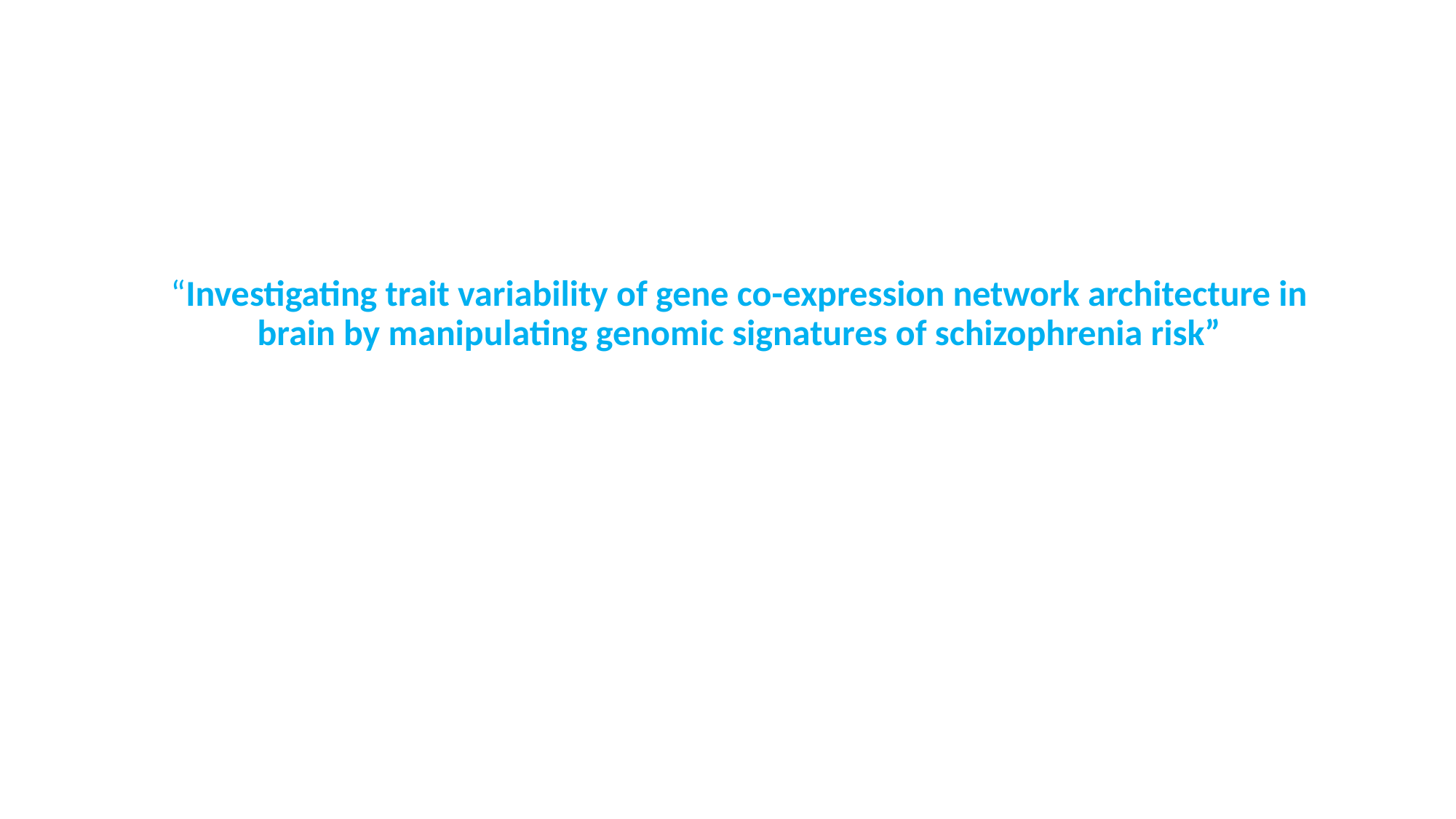

“Investigating trait variability of gene co-expression network architecture in brain by manipulating genomic signatures of schizophrenia risk”

### Slide 2
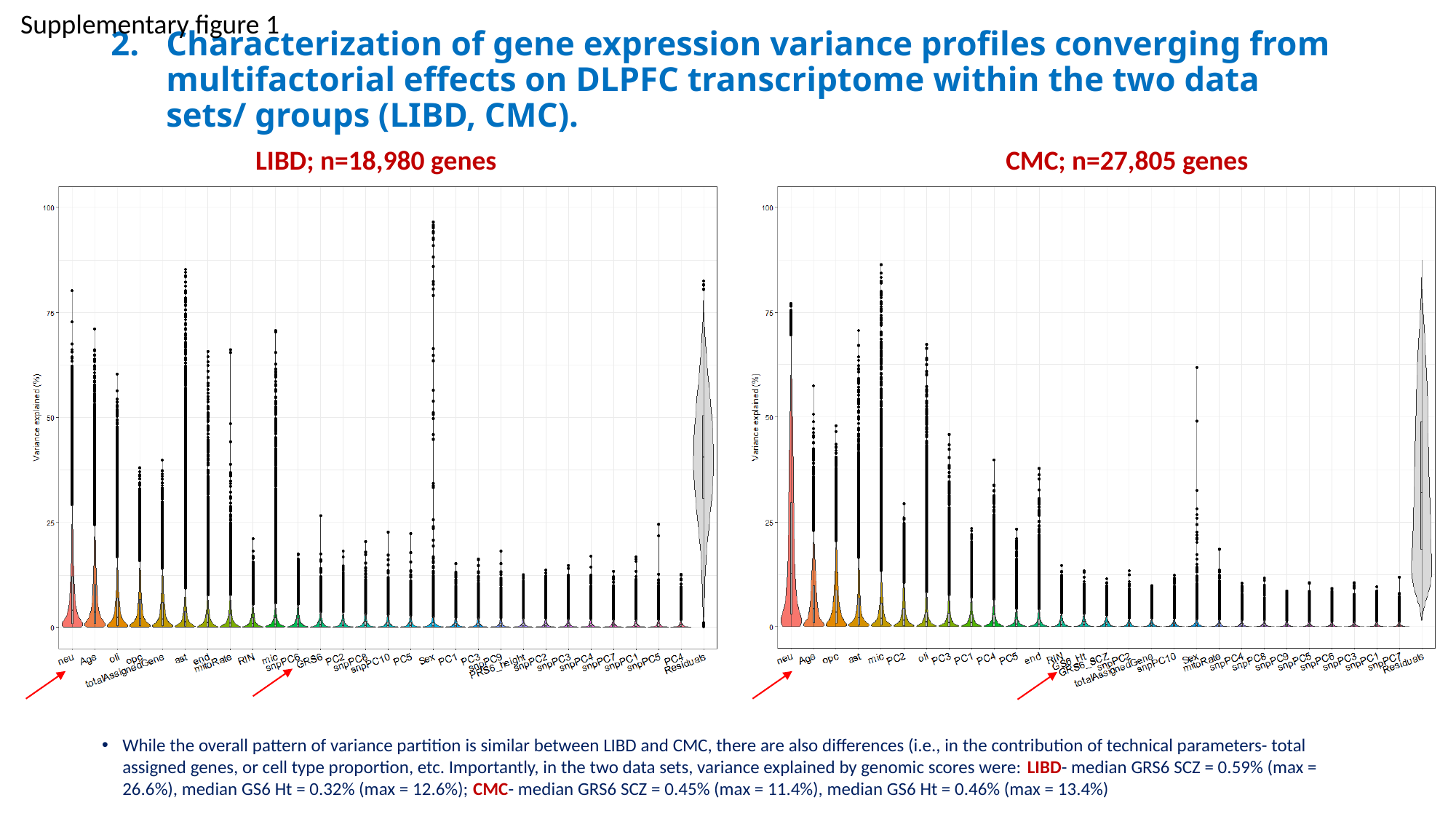

Supplementary figure 1
#### Characterization of gene expression variance profiles converging from multifactorial effects on DLPFC transcriptome within the two data sets/ groups (LIBD, CMC).
LIBD; n=18,980 genes
CMC; n=27,805 genes
While the overall pattern of variance partition is similar between LIBD and CMC, there are also differences (i.e., in the contribution of technical parameters- total assigned genes, or cell type proportion, etc. Importantly, in the two data sets, variance explained by genomic scores were: LIBD- median GRS6 SCZ = 0.59% (max = 26.6%), median GS6 Ht = 0.32% (max = 12.6%); CMC- median GRS6 SCZ = 0.45% (max = 11.4%), median GS6 Ht = 0.46% (max = 13.4%)

### Slide 3
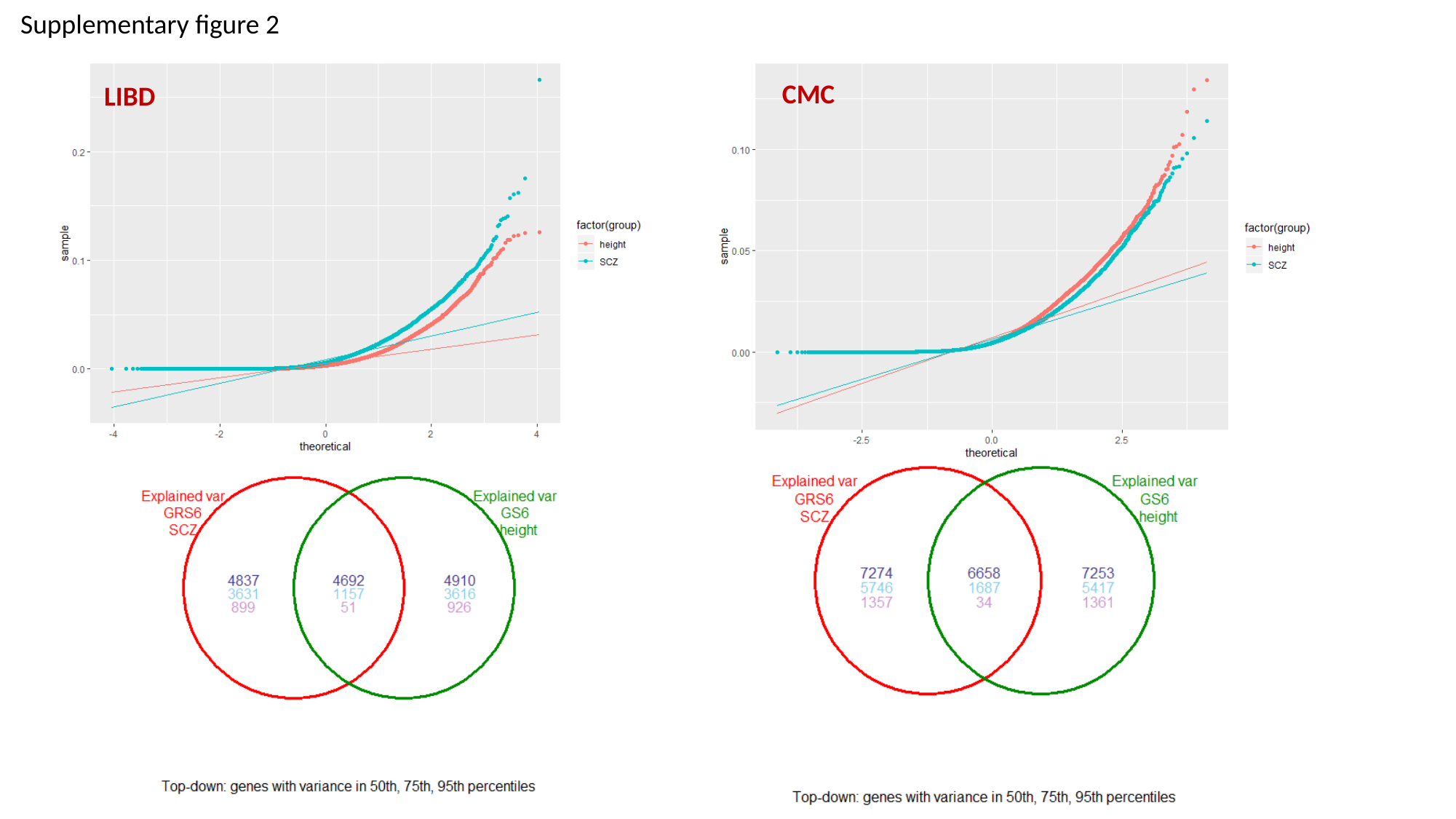

Supplementary figure 2
CMC
LIBD

### Slide 4
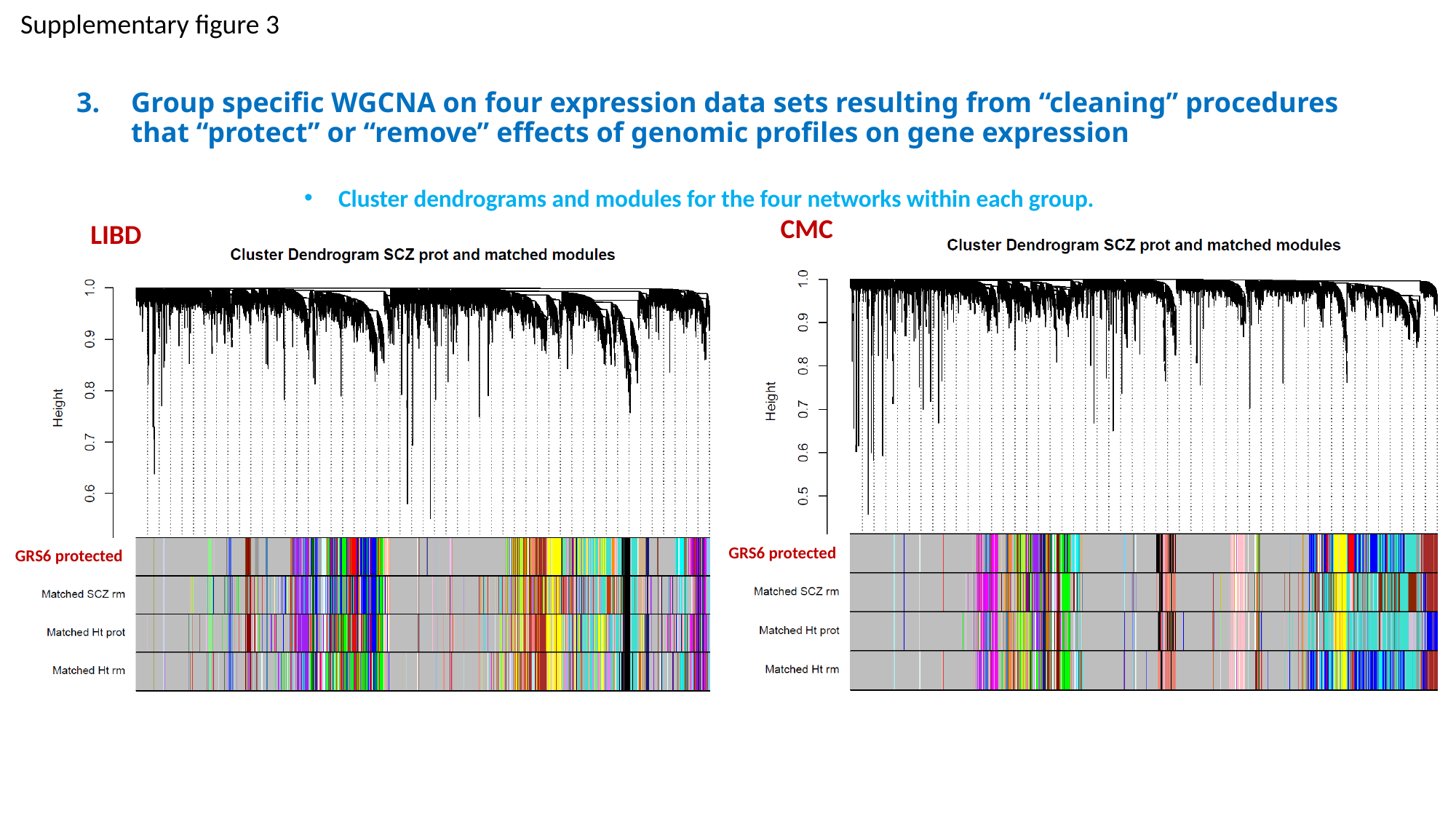

Supplementary figure 3
Group specific WGCNA on four expression data sets resulting from “cleaning” procedures that “protect” or “remove” effects of genomic profiles on gene expression
Cluster dendrograms and modules for the four networks within each group.
CMC
GRS6 protected
LIBD
GRS6 protected

### Slide 5
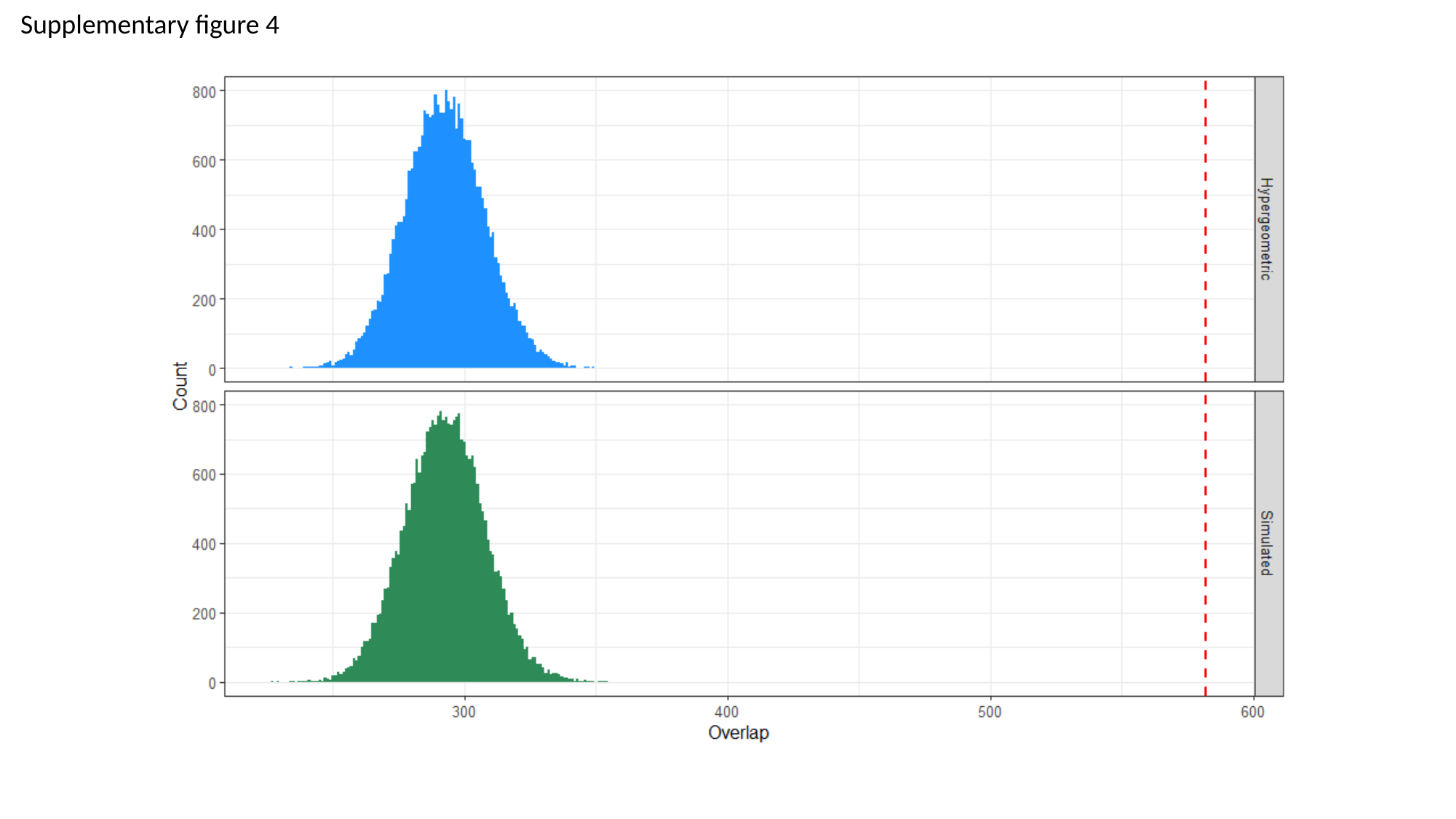

Supplementary figure 4

### Slide 6
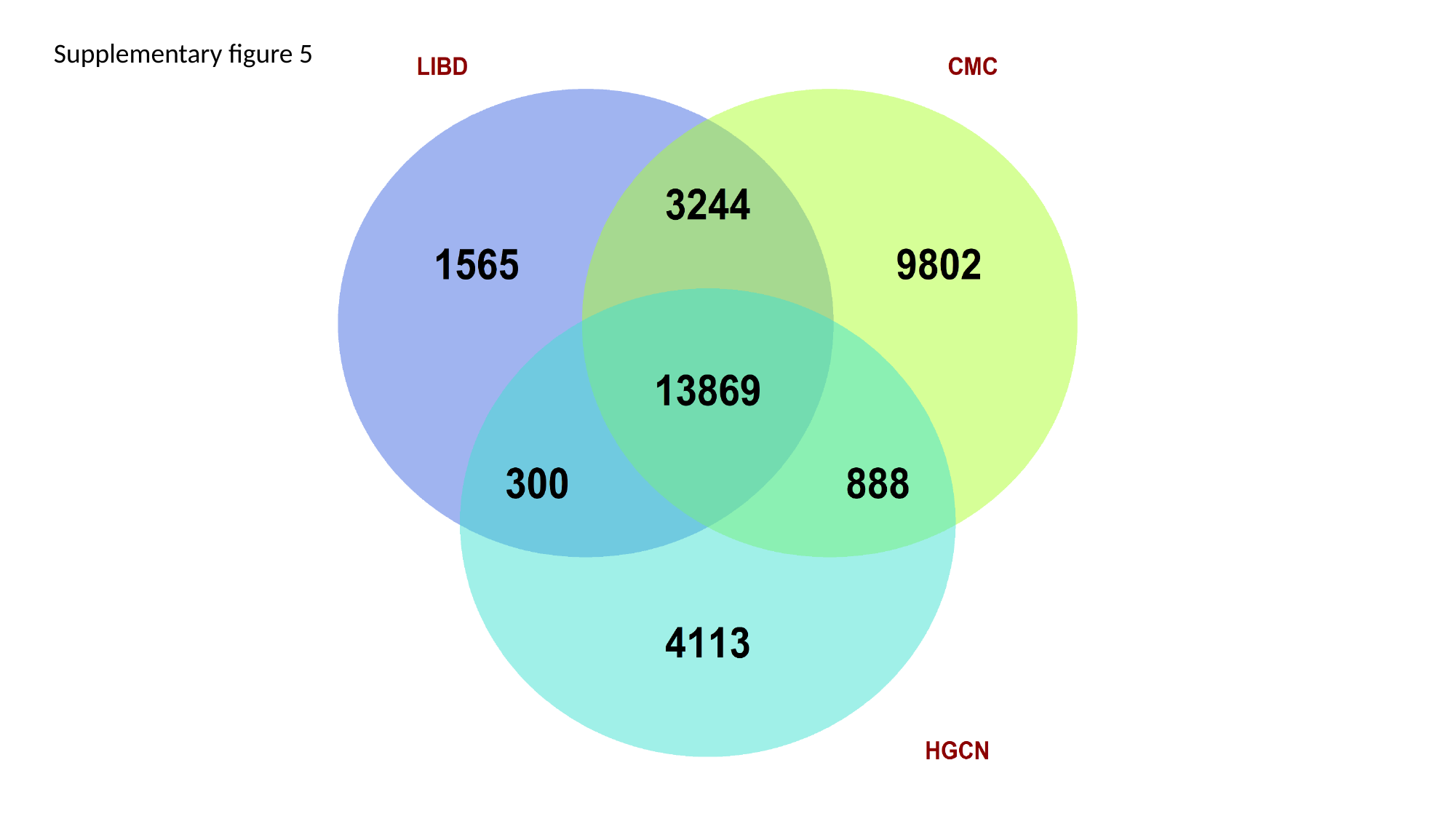

Supplementary figure 5
